# No evidence for motion dazzle in an evolutionary citizen science game

**DOI:** 10.1101/792614

**Authors:** Anna E. Hughes, David Griffiths, Jolyon Troscianko, Laura A. Kelley

## Abstract

The motion dazzle hypothesis posits that high contrast geometric patterns can cause difficulties in tracking a moving target, and has been argued to explain the patterning of animals such as zebras. Research to date has only tested a small number of patterns, offering equivocal support for the hypothesis. Here, we take a genetic programming approach to allow patterns to evolve based on their fitness (time taken to capture) and thus find the optimal strategy for providing protection when moving. Our ‘Dazzle Bug’ citizen science game tested over 1.5 million targets in a touch screen game at a popular visitor attraction. Surprisingly, we found that targets lost pattern elements during evolution and became closely background matching. Modelling results suggested that targets with lower motion energy were harder to catch. Our results indicate that low contrast, featureless targets offer the greatest protection against capture when in motion, challenging the motion dazzle hypothesis.

## Introduction

The high contrast, conspicuous patterns seen on animals such as zebra, snakes and fishes have attracted a range of evolutionary explanations, including camouflage, thermoregulation, communication and the avoidance of biting flies [1–7]. One hypothesis that has received attention in recent years is the ‘motion dazzle’ hypothesis, which proposes that these patterns may act to cause confusion when the animal is in motion, causing illusions in the visual system of the viewer that may lead to misjudgements of speed and direction[8].

There have been a number of studies that have provided support for the motion dazzle hypothesis. For example, it has been shown that putative dazzle patterns are relatively difficult for humans to ‘catch’ in a computer based touch screen game [9–11], and may also interfere with speed [12–14] and direction[15] perception. There is also evidence that some orientations of stripes can interfere with the ability to track one target within a larger group [16–18]. Finally, modelling work has suggested that striped patterns may be particularly prone to creating erroneous motion signals in the visual system, which may underlie these types of behavioural effects[19].

Despite these findings, not all research has supported the motion dazzle hypothesis. Some studies on humans have found that striped targets are easier to capture than non-patterned targets [20,21], and moving cuttlefish have been shown to preferentially display low contrast patterns[22]. Similarly, a recent study using natural predators hunting patterned prey found no evidence for a benefit of motion dazzle patterning compared to uniform coloration[23]. Even studies which have argued for an effect of motion dazzle patterning have normally shown that there is no benefit in terms of capture success of striped patterning over a luminance matched non-patterned target, suggesting that the benefit of stripes may not be unique [9,11,14,21].

One limitation of previous studies is that they have tested a relatively small range of patterns, often chosen arbitrarily. This means that it is not yet clear whether we have truly discovered the optimal patterning type to provide protection when in motion; it may be that there are more effective options than those tested so far. However, the small-scale psychophysics-style experiments used to date make it difficult to test large numbers of patterns. We therefore took a novel approach, using genetic programming to allow the patterning of targets to ‘evolve’ across generations in response to capture success [24–26]. In this way, we can ask which patterning strategy is optimal, given the almost infinite number of possible patterns that can be generated. To obtain the large amount of data required for this approach, we ran our experiment as a citizen science game (‘Dazzle Bug’) in a popular visitor attraction. Participants played the game by tapping on the moving targets (‘bugs’) with their finger as quickly as possible in order to ‘catch’ them (Figure 1). We ran a number of replicates of the evolutionary process for three populations of different speeds, to assess whether the optimal patterning changes as a function of the target movement speed.

**Figure 1:**
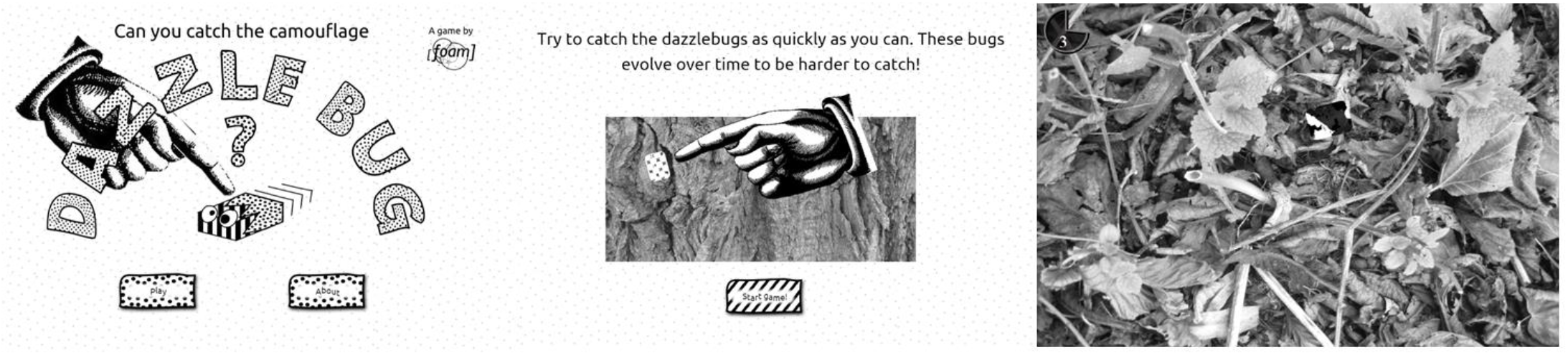
Figure showing screenshots from the game. Left: title screen. Middle: instructions presented to the participant. Right: the game in progress. Participants could see the time remaining on the trial via the countdown clock in the top left hand corner.

Our first aim was to demonstrate a fitness increase in our experimental populations, which we defined as an increase in the average capture time across generations. We did this by comparing to a simulation run of the evolutionary algorithm, using randomised capture times. We then investigated how the target patterning changed across generations for different speed populations, using image analysis to measure contrast and the presence of stripes at different orientations. We also looked at whether selection rates differed for the different speed populations, using the Land, Arnold and Wade framework [27–29], allowing us to consider how selection pressure might vary across the generations. Finally, we asked whether motion perception modelling can help to explain our experimental results.

## Results

### Is there a fitness increase for the experimental populations, and does this differ from the null population?

Figure 2 shows there were clearly large differences in fitness (capture speed) among populations, with the fast bugs being hardest to catch, followed by the medium bugs and then finally the slow bugs (χ^2^ = 50892.85, p < 0.001). There was a considerable level of noise in the data, which is to be expected given the wide range of participants and fast reactions required. Nevertheless, there was also a significant increase in fitness across generations (χ^2^ = 208.72, p < 0.001). Increases were often particularly obvious in the early generations of the game.

**Figure 2:**
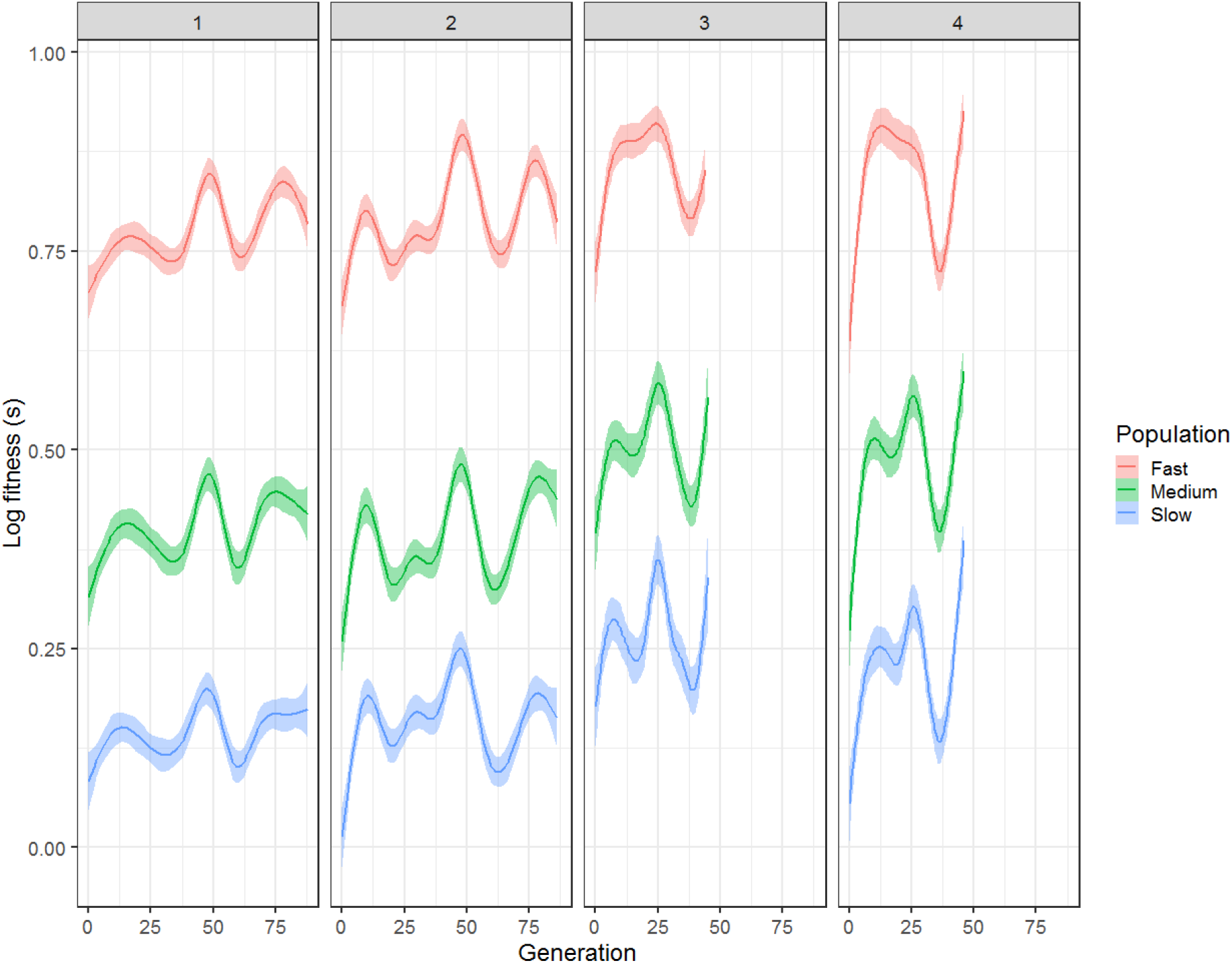
Experimental data (shown as a smoothed GAM) from all four replicates, showing how log fitness changes as a function of generation number and speed population.

The experimental data also show a significant difference in fitness change compared to the null data (interaction between dataset and second order effect of generation: χ^2^ = 161.985, p < 0.001). The experimental data shows a characteristic quadratic shape, with an initial increase that flattens off (Figure 3). We therefore have evidence for a fitness increase in our experimental population, suggesting that selection is occurring to optimise patterning types.

**Figure 3:**
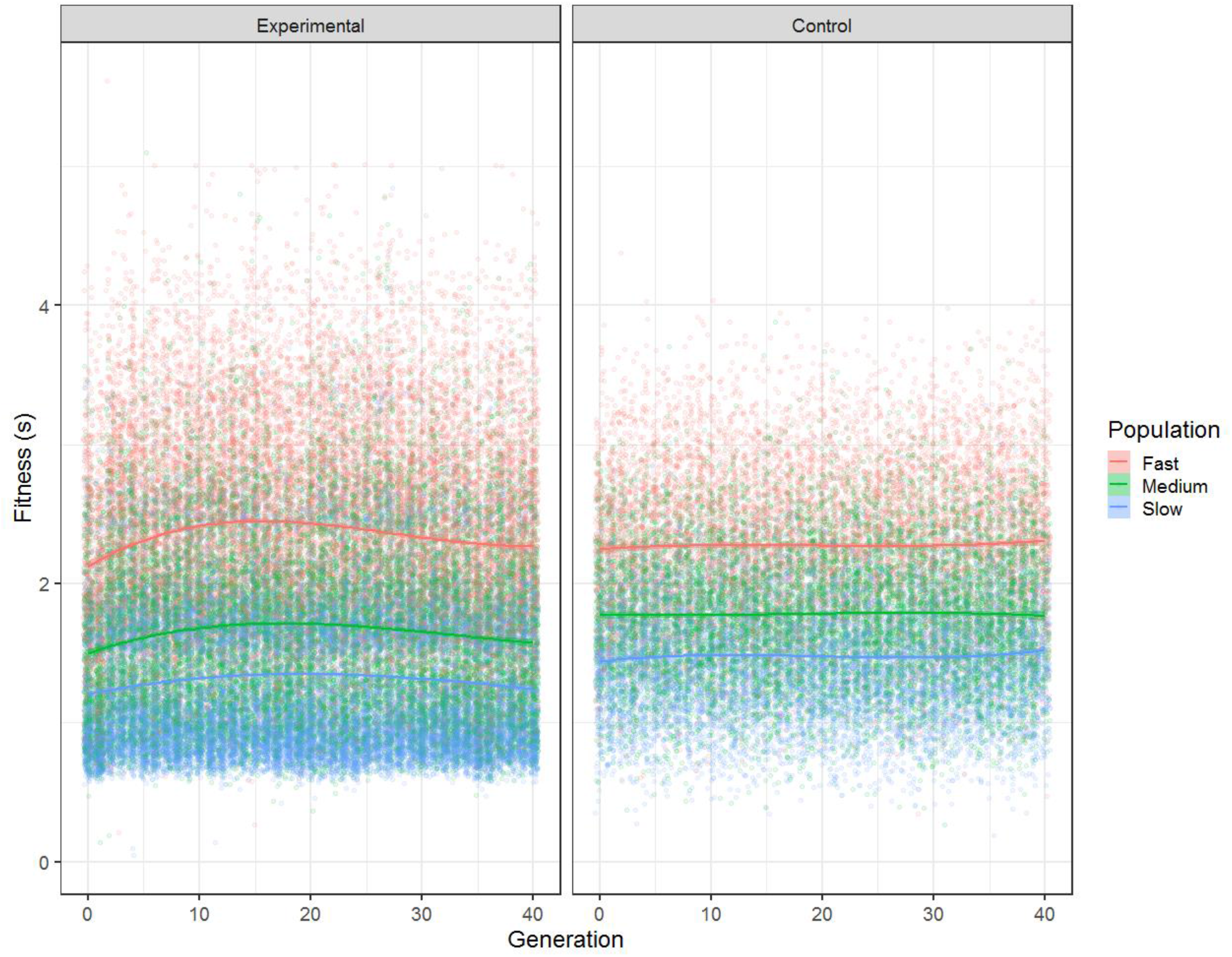
Experimental data (left) and control data (right) compared across 40 generations and for the three different speed populations. Experimental data has been collapsed across all 4 replicates. All raw data points are plotted and the curves are fit using splines with two degrees of freedom.

### How does bug patterning change in the experimental and null populations?

All four populations of evolving bugs demonstrated a loss of pattern information over the generations – converging on uniform background-matching colours (Figure 4, top) – while the control populations maintained their pattern diversity (Figure 4, bottom). Quantifying this using our five most informative pattern metrics (Figure 5) shows that there are always clear differences between how the pattern metrics change in the experimental condition compared to the control condition (interaction between experimental/control condition and pattern metric for cumulative link models - standard deviation of bug luminance: χ^2^ = 36207.5, p < 0.001; vertical stripes: χ^2^ = 36848.3, p < 0.001; horizontal stripes: χ^2^ = 36587.0, p < 0.001; diagonal stripes: χ^2^ = 36299.0, p < 0.001; right edge: χ^2^ = 36613.3, p < 0.001). Broadly, there always seems to be an overall decrease in pattern complexity in the experimental case, whereas there is much more variability in the control condition.

**Figure 4:**
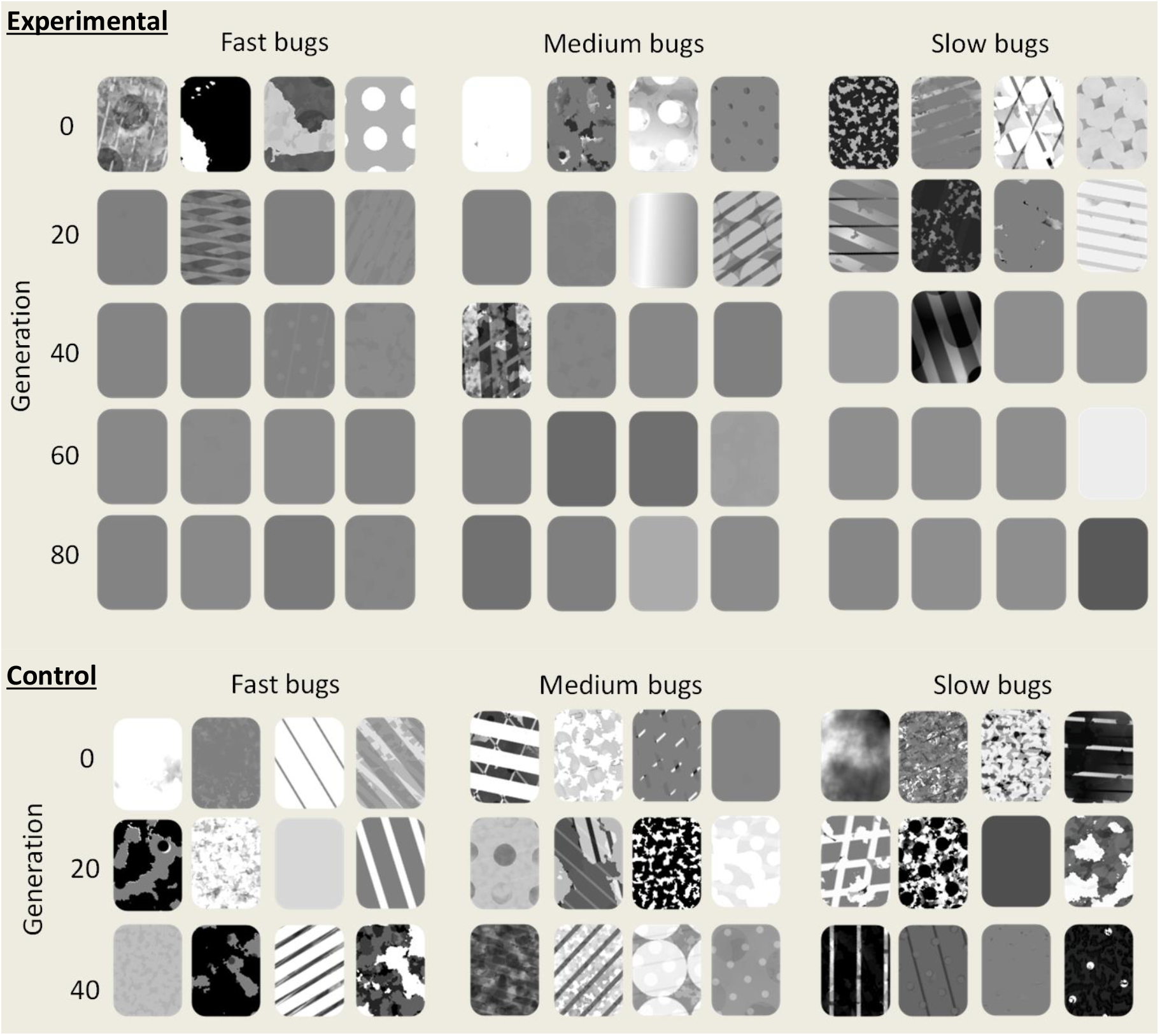
Top - random bugs from generations 0, 20, 40, 60 and 80 (all from the same replicate) of the experimental data, split into populations (fast, medium and slow). Bottom- random bugs from generations 0, 20, and 40 (all from the same replicate) of the control data, split into populations (fast, medium and slow).

**Figure 5:**
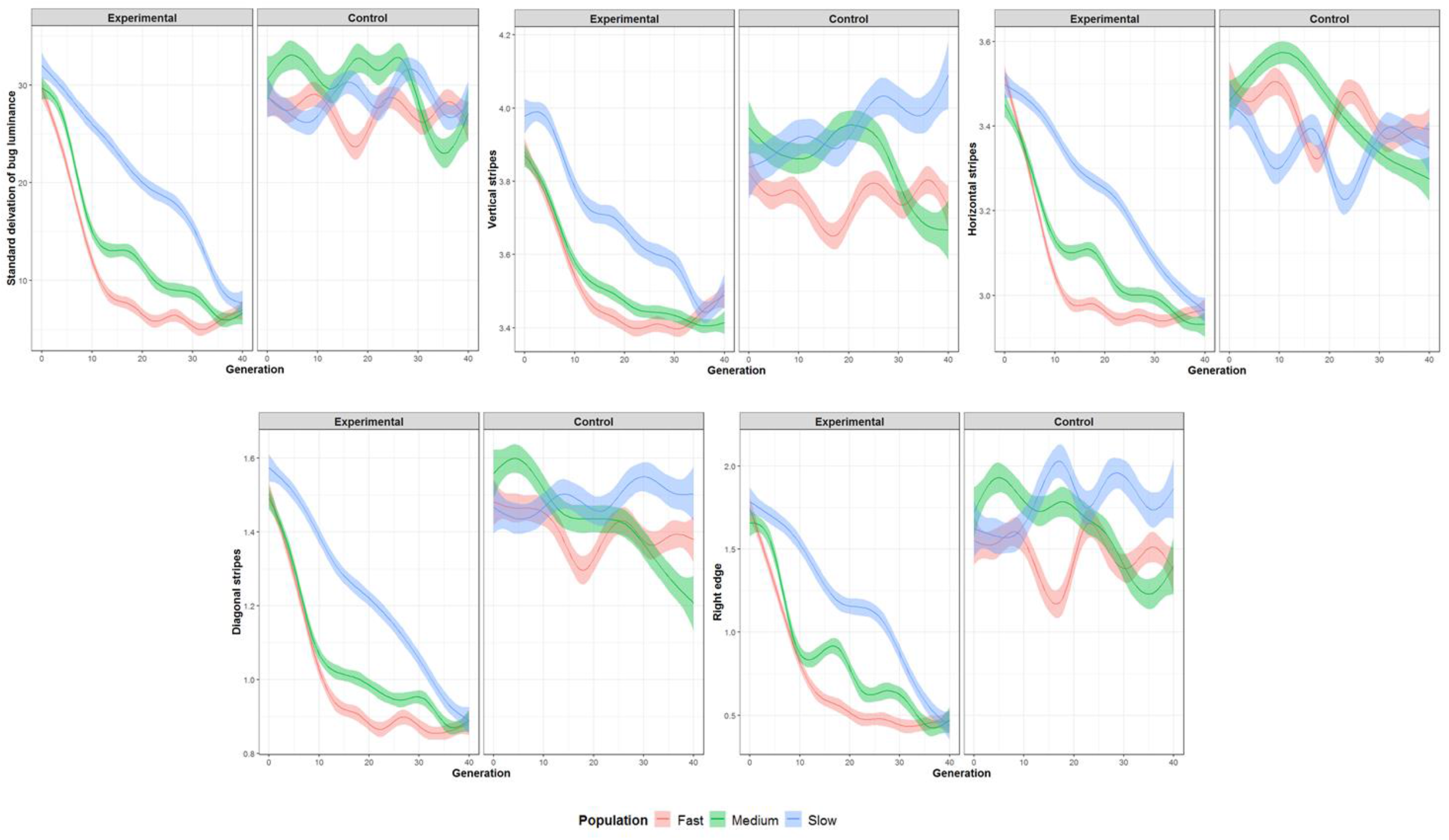
Change in parameters across generations, both for the experimental (left) and control (right) conditions, for five pattern metrics: from top left, these are the standard deviation of the bug luminance, the pattern energy for the vertical stripes, horizontal stripes, diagonal stripes and the right edge.

### Are there differences in selection rate for each speed population?

The data allow us to determine the main selection pressures operating on each population of bugs within each generation (normalised linear selection rates (β)), so that we can assess whether pressures change over evolutionary time. Differences in selection rates across generations were seen for luminance (χ^2^ = 12.815, p = 0.002), vertical stripes (χ^2^ = 11.593, p = 0.003) and for diagonal stripes (χ^2^ = 6.647, p = 0.036). There was no evidence for difference in selection rates for both the horizontal stripe (χ^2^ = 1.705, p = 0.426) and the right edge metrics (χ^2^ = 5.486, p = 0.064).

The standard deviation of the luminance of the bugs appears to be particularly important for the ‘fast’ population; there is strong selection pressure particularly in early generations, and this differs from the selection rate seen in the ‘medium’ and ‘slow’ populations (Figure 6; fast-medium comparison: t = −2.883, p = 0.012; fast-slow comparison: t = −3.138, p = 0.005; medium-slow comparison: t = −0.249, p = 0.967). For vertical stripes, there is some evidence for stronger selection pressure for medium compared to slow bugs (t = −2.654, p = 0.022). For all other patterning parameters, there were no significant differences between the different speed populations (horizontal stripes − χ^2^ = 3.928, p = 0.140, diagonal stripes − χ^2^ = 1.783, p = 0.410, right edge − χ^2^ =4.330, p = 0.115).

**Figure 6:**
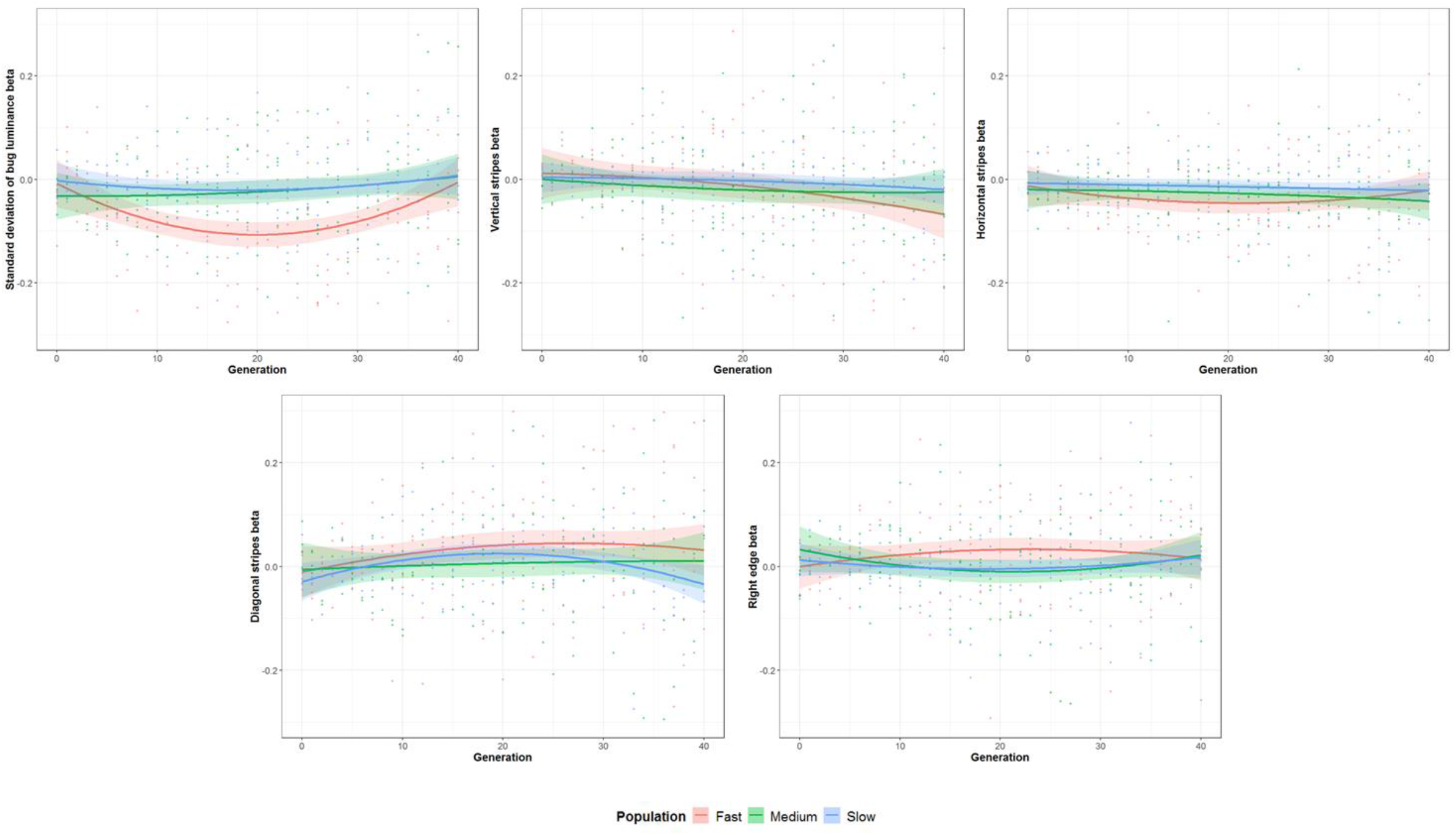
Normalised linear selection rates (β) for the five most important camouflage metrics across generations. Polynomial curves are fitted to the raw data up to generation 40. Individual points show the selection rates for each replicate.

### Can motion modelling help to explain the experimental findings?

According to previous modelling work[19], we would expect targets to produce strong motion illusions if they are both highly coherent (the motion vectors produced tend to be in a highly similar direction) and biased (the average trajectory of the motion vectors is quite different from the ‘veridical’ direction of the target). When considering only the most coherent targets, there is a significant relationship between bias and fitness (Figure 7); the fitness of the targets increased as the bias increased (interaction between coherence and bias: F = 5.985, p = 0.015), in line with previous predictions[19]. In addition, the targets with the highest bias also tended to be relatively stripy and high contrast (bugs with higher bias had both higher standard deviations of luminance F = 10.844, p =0.001, and levels of vertical stripes F = 35.688, p < 0.001) again suggesting that these “motion dazzle” type patterns might be expected to create illusory motion signals.

**Figure 7:**
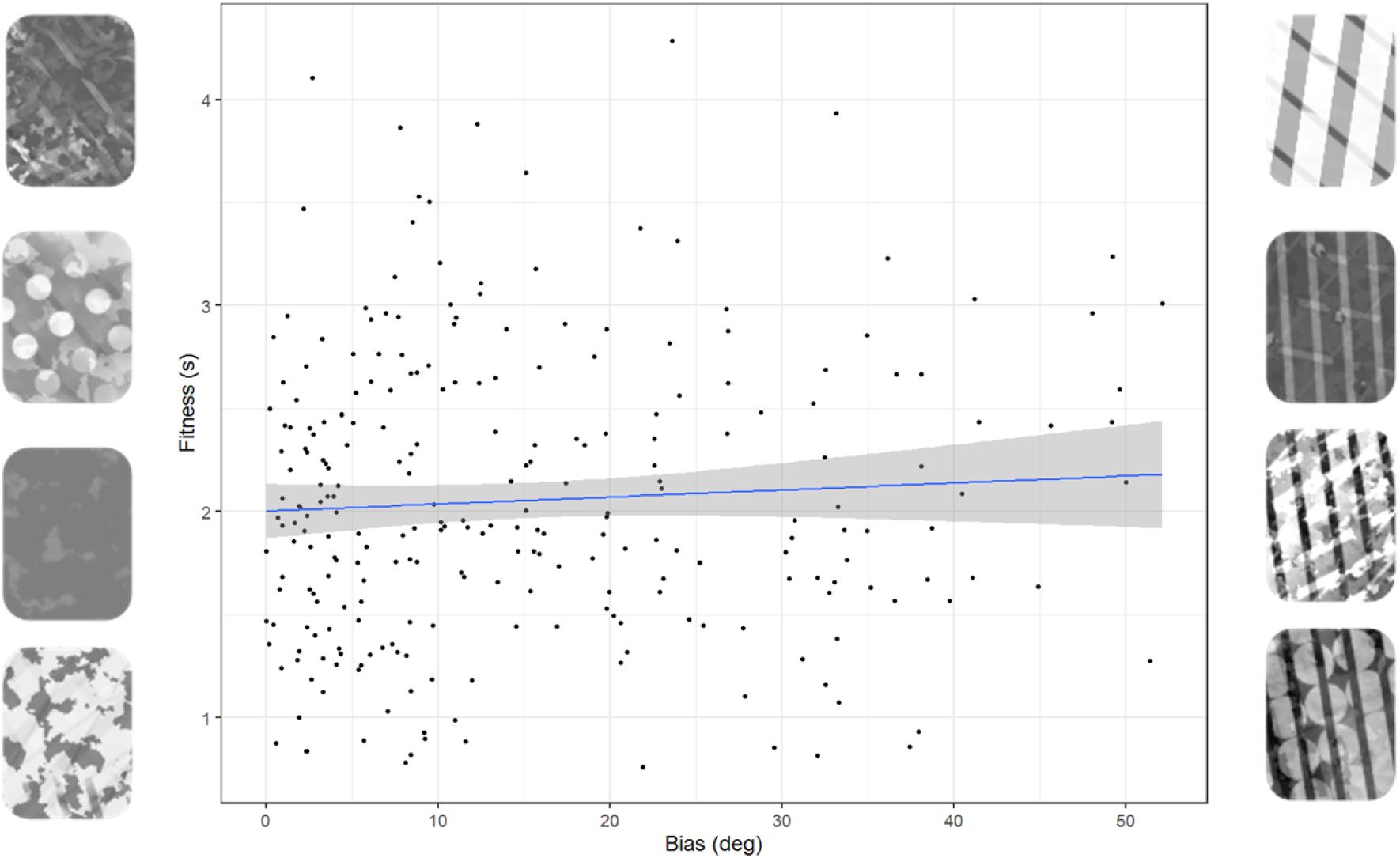
Data from the generation 0 fast bugs, using only those bugs above the median coherence value (i.e. relatively highly coherent targets) and plotting fitness against the bias. Exemplars on the left are bugs that had low bias values, according to the motion model; exemplars on the right are bugs that had high bias values.

However, these results do not seem to explain our evolutionary findings, where we saw a strong tendency for targets to become lower contrast and non-patterned. A second metric from our motion modelling is the motion energy, which can be conceptualised as how salient or visible the motion is. Here, there is a very different relationship with fitness, as can be seen in Figure 8, with low motion energy targets (that tend to be low contrast and have little patterning) having higher fitness than those with higher motion energy (that tend to have high contrast and strong patterning) (F = 4.391, p = 0.027; F = 4.989, p = 0.026 if data were not filtered to exclude cases with a circular mean difference of greater than 6 degrees). Bugs with higher mean vector lengths had both higher standard deviations of luminance (F = 1171.8, p < 0.001) and levels of vertical striping (F = 545, p < 0.001).

**Figure 8:**
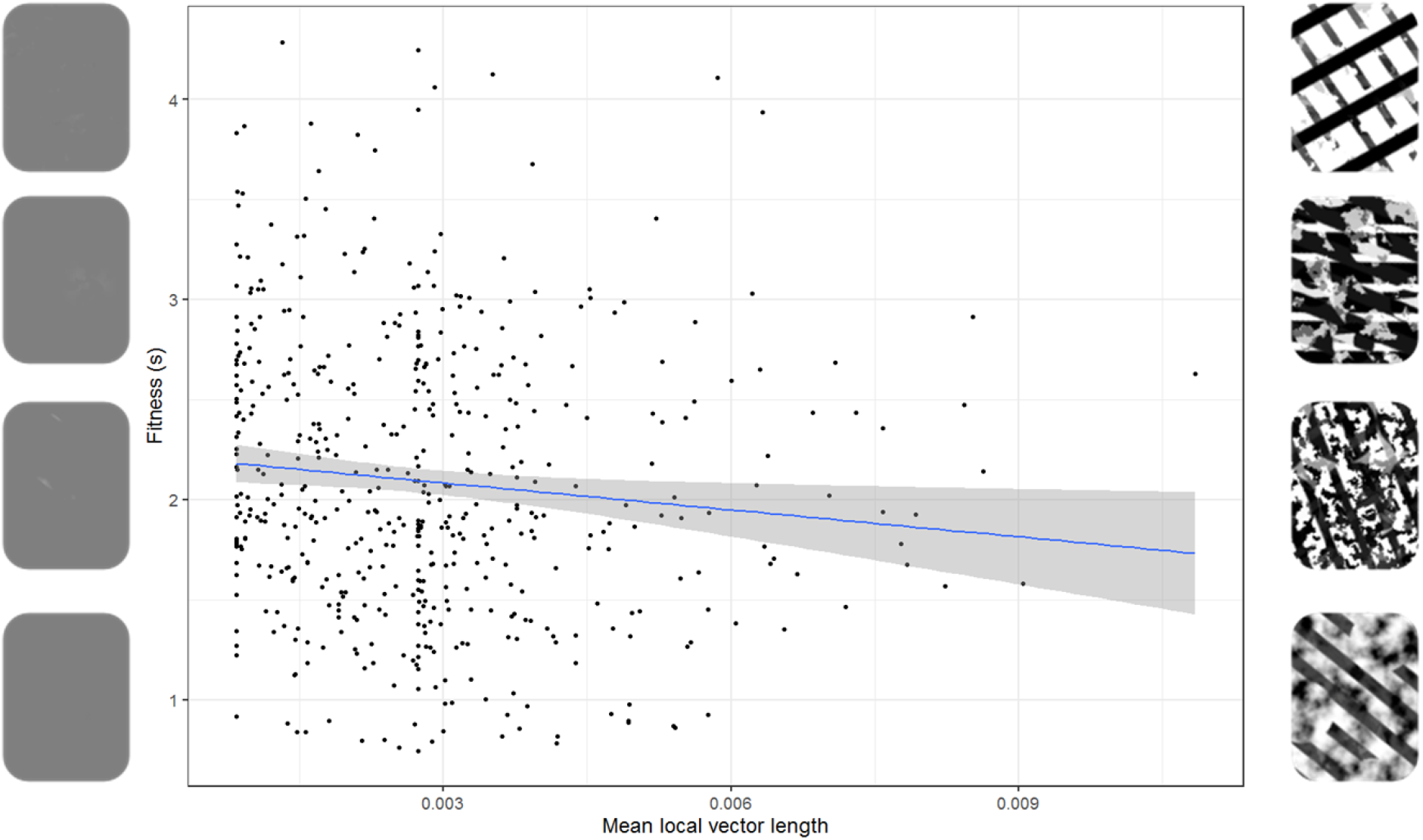
Data from the generation 0 fast bugs, plotting fitness against the mean local vector length (a measure of motion energy). Exemplars on the left are bugs that had low motion energy values, according to the motion model; exemplars on the right are bugs that had high motion energy values.

## Discussion

Using a large-scale evolutionary citizen science game, we found no evidence that putative ‘motion dazzle’ patterning can offer protection when in motion; despite predictions that high contrast, geometric patterning should cause visual illusions that make targets harder to catch, we found that the targets consistently evolved to become less patterned and lower contrast. This happened for all speeds tested and all replicates of the experiment, although these changes seemed to occur more quickly in populations with faster speeds. Motion modelling suggested that these results could be a consequence of the motion energy of the stimulus, as this correlated with capture time, with lower motion energy targets being more difficult to catch. Our results have important consequences for our understanding of the evolution of stripes, and for how animals should best protect themselves from capture when in motion.

Our results are perhaps surprising in the context of most literature on motion dazzle to date, which has suggested that stripes seem to be relatively difficult to catch or can cause illusions of speed or direction perception [9–12,14–18]. However, we note that there has indeed been plenty of evidence in the literature for uniform grey patterns also being relatively difficult to catch, and in some cases perhaps even harder than striped targets. For example, grey targets always survive well in capture studies [9,11,14,21]. Similarly, in tracking tasks, low contrast parallel stripes were found to be more difficult to track than high contrast parallel stripes[18], arguing against a motion dazzle explanation. Recent work has also suggested that in some cases striped patterns are only difficult to catch when the targets are moving sufficiently quickly to blend via the “flicker-fusion” effect into uniform grey[43]. Our results therefore suggest that uniform grey targets had a survival advantage over other types of target patterning, leading them to become fixed as the optimal strategy in all our populations, regardless of speed or replicate number.

Motion modelling has previously suggested that stripes should create erroneous motion signals that are both highly coherent and biased[19], implying striped prey should be more difficult to catch. However, to our knowledge, modelling results have not previously been compared to behavioural data. Our large dataset therefore offers a perfect opportunity to study whether the motion modelling results do indeed correlate with capture times. In support of the motion dazzle hypothesis[19], we do indeed find that highly coherent and biased targets tend to be more difficult to catch than less biased coherent targets, and that the most biased and coherent targets are often stripy. However, this clearly does not explain the results we see in the evolutionary game. We thus considered another metric that can be calculated from motion models, namely the motion energy, and found that this also correlated with capture success. Targets with low motion energy (that tended to be uniform grey) were harder to catch than targets with high motion energy (that were much more high contrast and patterned).

Why does background-matching (reducing motion energy) seem to be a better predictor of the outcomes in our evolutionary games compared to motion dazzle strategies which maximise the bias/coherence metric? We speculate that motion energy is a very consistent signal; regardless of the trajectory of the bug or the speed, the targets with low visibility will be harder to catch than those that are highly visible. We propose that the effects of stripes may be much more dependent on the particular orientation of the stripes, given that the most effective striped targets appeared to have relatively similar dominant orientations (Figure 7), and previous studies have shown orientation dependence for the effects of striped targets [16–18,21]. Small mutations affecting the rotation of striped patterns could therefore potentially cause large changes in fitness, potentially making striped patterns a relatively unstable evolutionary strategy compared to uniform grey in our experiment. This could suggest that other factors may play a role in maintaining striped patterning in animals, and it would be instructive for future studies to more closely consider the possibility of stripes serving multiple functions.

We used three different speed populations in order to assess whether there were differences in the patterns that evolved. As expected, we found that there were strong differences in capture difficulty for different speed populations, with fast targets being the hardest to capture, but we did not find evidence for there being differences in the target patterning that evolved, with all populations becoming uniform grey. This is in agreement with previous work suggesting that there is no interaction between target speed and prey patterning[11], at least for speeds below that needed to create a “flicker-fusion” effect. However, we did find increased selection in “fast” populations, particularly early on in the evolutionary process for the contrast metric and later on for the vertical stripe metric. This may simply reflect the higher difficulty of these targets, which is likely to give a wider range of capture times and thus offer more variation for selection to operate on, potentially exaggerating the selection process.

Genetic algorithms are complex and there are many different ways to implement them [24–26]. We therefore carried out control experiments using simulated reaction time data with similar average distributions to the real data, helping us to rule out explanations of our results based on algorithmic biases or genetic drift. Our results show clearly that selection pressures do indeed operate in our game and that the change towards grey targets does not simply reflect drift. However, while our set-up allowed us to explore a very wide range of pattern types, it is possible that different algorithms could produce different targets and thus perhaps different results. For example, our targets were rarely highly asymmetric (although this was possible). Recent research has suggested that stripes may be particularly effective at misdirecting capture attempts when they are placed on the anterior of a target[13], suggesting that an interesting direction for future work could be to allow the algorithm to specify different genes (and thus different patterning) for different parts of the target.

Our experiment used human participants, in line with the majority of studies in this area. Of course, in the natural world, the viewing animals might have very different visual systems to humans. We removed colour cues from our experiment, as it is well known that different species have very different colour perception [44,45], although motion vision is generally thought to be predominantly achromatic [46–48]. However, there is also large variability in the perception of temporal changes across different species[49] which we could not adequately compensate for in this experimental set up. Despite this, our main conclusions broadly agree with previous studies carried out on non-human predators and prey [22,23]. However, it would of course be highly instructive to carry out similar experiments with non-human animal participants to determine whether the results we report here are more widely generalizable.

Overall, we find limited evidence for motion dazzle effects in a citizen science evolutionary game, which we believe is the most comprehensive test of this hypothesis to date. Stripes were able to cause motion illusions and reduce capture times in some scenarios, meaning that there may still be specific cases where motion dazzle can be at least part of an explanation for the evolution of striped patterns. However, our results suggest that uniform grey targets appear to be a more stable optimal solution.

## Methods

### Subjects

We did not collect any demographic data from participants. This was to streamline participation in the study (which was conducted in a busy exhibition space) and also because it would be difficult to verify the accuracy of the information presented. To overcome the limitations of being unable to account for participant age, handedness and gender, we collected a large sample size of participants over many generations (1,554,935 targets were caught in total across the whole experiment, involving approximately 75,000 participants). This project was carried out with ethical approval from the University of Cambridge (pre.2014.08).

### Experimental methods

The Dazzle Bug game was installed at the Eden Project (St. Austell, UK) on a touch screen computer as part of an interactive exhibition, and the data used were collected between May 2018 and January 2019. The game was coded in HTML5 canvas (source code is available at https://github.com/nebogeo/dazzlebug, DOI: 10.5281/zenodo.2560935) and is playable online at dazzle-bug.co.uk/exhib.html (the online data are not analysed in this paper). The screen had an area of 478 × 269mm and the screen resolution of the game was 1237 × 820 pixels. The viewing distance of participants to the screen was approximately 60cm.

The game had a similar format to many previous studies testing motion dazzle effects [10,11,21] in that participants were presented with a small rectangular target (75 × 100 pixels, or 29.0 × 38.6mm; visual angle 2.76×3.69°) which they had to try to ‘catch’ as quickly as possible after it had appeared by touching it with their finger (Figure 1). Targets began their movement at a random position on the screen and moved with a linear trajectory. The angle of movement changed throughout a trial, both at the edge of the target arena via reflection (to ensure that the target remained visible to the participant) and randomly throughout movement (once every half a second, and when an unsuccessful capture attempt was made; the new angle was randomly chosen based on its previous angle plus or minus 90 degrees). Targets could be presented at one of three speeds, fast, medium or slow (600, 450 or 300 pixels per second respectively, independent of frame rate, which equated to 231.8, 173.8 and 115.9mm/s), and each participant was presented with a random mix of targets of all three speeds. Participants had 5 seconds to catch each target. After the target had been caught, or the time-out limit had been reached, the game would move automatically onto the next target. A game consisted of 20 trials in total, with the targets presented randomly selected from the current generation.

### Background photos

Targets were presented against one of 40 naturalistic background photographs (of e.g. grass, tree bark or leaf litter). The background was randomly selected on each trial. The photos were calibrated and converted to greyscale (with an average pixel value of 127).

### Pattern generation

The patterns throughout the game were generated through a genetic programming approach [24–26]. This does not attempt to directly mimic biological evolution, but is instead a method allowing the exploration of an unbounded parameter space in an efficient manner, using algorithms inspired by natural selection processes. The key principle is that the evolutionary process acts to modify small ‘computer programs’ that specify the patterning presented on each target. This allows a great deal of flexibility in the complexity of target patterning and reduces artificial bounds on the evolutionary space that can be introduced in more traditional genetic algorithm methods[25].

Targets were generated in a hierarchical manner, as shown in Figure 9. The ‘tree structure’ of the program determining the target pattering is composed of two different types of node. One type of node is the ‘terminal node’ that is found on the outer ring of the tree. There were two possible variants of terminal node (each chosen with a probability of 50%). One variant was a flat image of a specific RGB colour (always greyscale) and alpha (transparency) value. The second variant of terminal node consisted of a specific pre-generated image; there were 66 different initial images from a range of different categories, including striped patterns, spotted patterns and noise patterns, and with a range of spatial scales (see Figure 10). These base images could also be moved using an x− offset and a y-offset value (with the patterns wrapping around the target) and rotated (in radians). The other type of node was the ‘combination node’. Here, two image inputs were combined using one of the following randomly selected nodes:

- Source-over: the second image was drawn on top of the first image
- Source-atop: the second image was drawn only when it overlapped the first image (i.e. the second image was not drawn on the transparent parts of the first image)
- Destination-over: the second image was drawn behind the first image
- Lighter: when both images overlapped, the new colour was determined by adding the colour values
- Xor: the new image was made transparent when both the first and second images overlapped, and was drawn normally everywhere else

**Figure 9:**
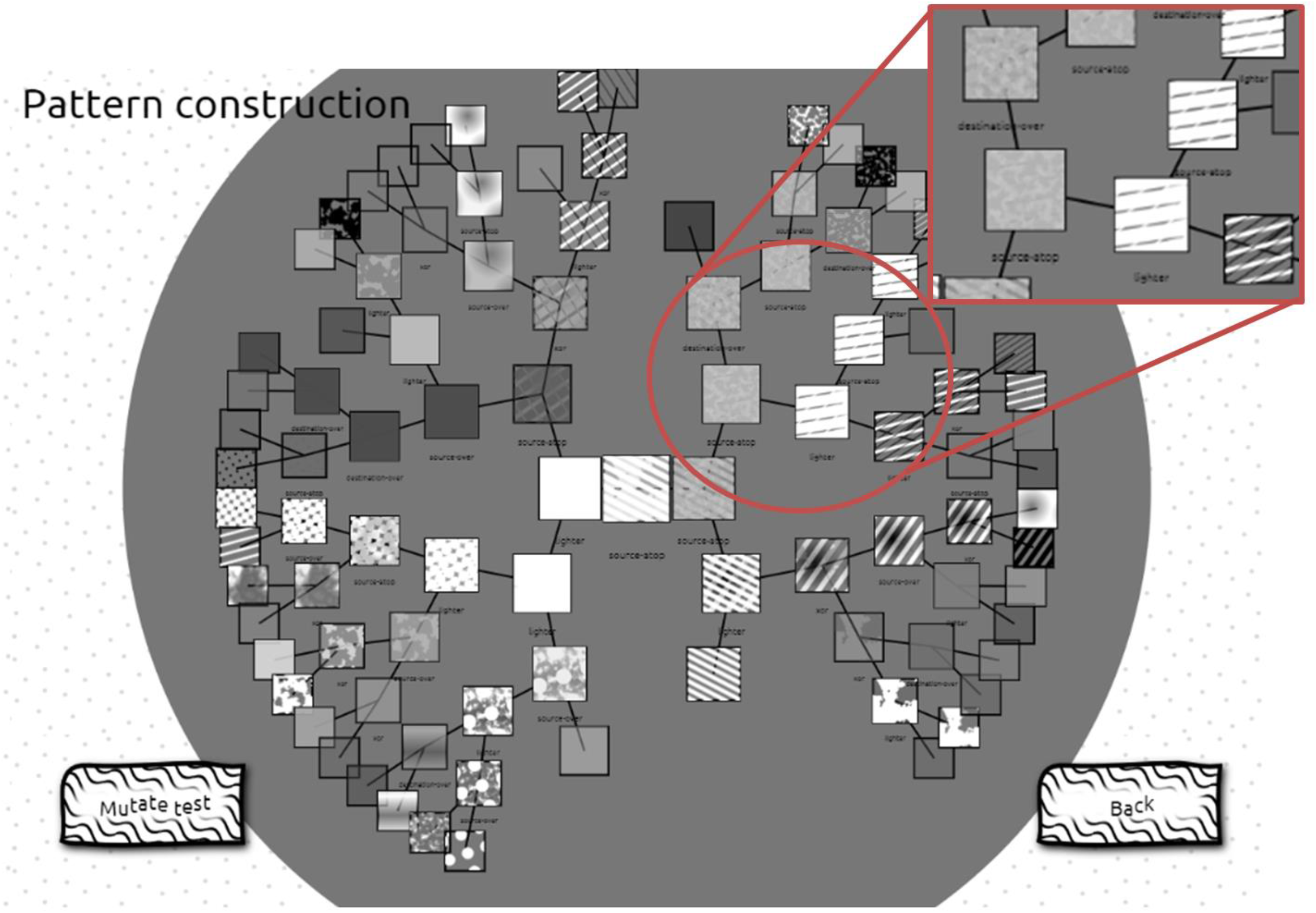
Schematic to show how targets are generated (available to view on the online version of the game). Each target can be thought of as being the top point of a ‘tree’ made up from a range of different images, combined in different ways. The ‘nodes’ of the tree are combination operations, that each take two images as input. The tree continues until the outer edge of the circle, where the terminal nodes are made up of the base images. The magnified region shows the combination operations more clearly.

**Figure 10:**
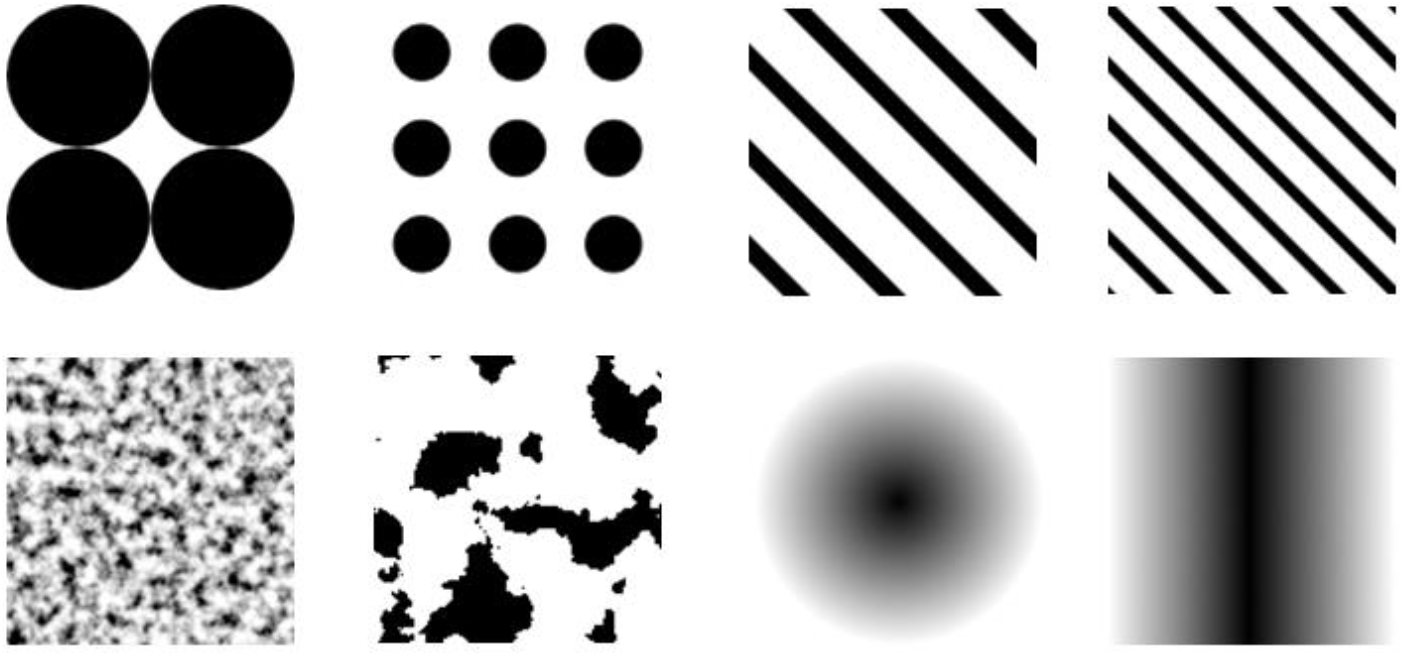
Selection of example base pre-generated images, including spots, stripes, noise and gradients.

An example displayed target is shown in the centre of the screen in Figure 9, and was formed by the top combination node of the tree. The input to this combination node could either be other combination nodes (as seen in this example) or could include a terminal node as well (with 20% probability). The process can be followed backwards until the input to a combination node is two terminal nodes (with randomly chosen parameter inputs), ending that part of the ‘tree’ and forming an outer edge of base images.

### Evolutionary process

Four replicates of the game were run, with each replicate containing three separate populations for each speed (fast, medium and slow) that each evolved separately. The first generation of each population contained 128 individuals that were completely randomly generated in accordance with the pattern generation process detailed above. These were then presented to players randomly until they had all been played five times. At this point, each one was scored by averaging the time taken to catch them, and the bottom half of the generation based on this measure of fitness was removed from the population. (Normalisation of participant times was not possible due to the design of the evolutionary algorithm). The top 64 targets were copied with no mutation to form one half of the new generation, and then copied again with mutation to form the other half. The mutation process involved either random changes of a parameter variable (e.g. changing the RGB colour) or selecting a random part of the tree (either a combination node or a terminal node), copying it and pasting it onto another random part of the tree. Pruning then occurred if the mutation process increased the depth of the tree to beyond the maximum permitted (6 layers). This process could lead to both increases and decreases in target complexity. The mutation rate was randomly selected for each target, with there being a 0-10% chance of a mutation occurring, but with the probability being weighted towards 0% (i.e. no mutation was most likely, but up to a 10% chance was possible).

The exact number of generations tested varied between replicates because each participant was randomly assigned to one replicate, and because not all replicates were run simultaneously. Replicate 1 had 89 generations, replicate 2 had 87 generations, replicate 3 had 45 generations and replicate 4 had 46 generations.

### Control model

We ran a control model to confirm that any systematic patterning changes seen during the real game were due to directional selection, rather than drift or biases within the genetic programming algorithm. This was set up identically to the real experiment, except that instead of participants playing the game, the computer randomly selected a ‘capture time’ for each target in each generation, based on a Gaussian distribution using the mean and standard deviation of each population in the real experiment (as individual clicks were not recorded in our experimental data, we estimated the variance of individual plays by multiplying the variance of the ‘bug-level’ fitness by the number of plays of each bug e.g. by 5). The null model was run for 40 generations.

### Quantification and statistical analysis

We analysed the patterning of the targets using custom written scripts in ImageJ (version 1.51k) [30]. This script first calculated the mean, minimum and maximum luminance of each target, and the standard deviation of the luminance. We also calculated the contrast of the target as the coefficient of variance in luminance (the standard deviation divided by the mean). We then used Gabor filtering methods that allow measurement of different angles at different spatial frequencies to determine the strength of these signals on the targets in a biologically plausible way [31–33]. We analysed four angles (vertical, horizontal, and two diagonal stripes) each at four different spatial frequencies (sigma values of 2, 4, 8 and 18 pixels). For each of these conditions, we calculated the standard deviation of Gabor-convolved pixel values as a measure of the “energy” at that particular angle and spatial frequency. Finally, we also measured the standard deviation of Gabor-convolved pixel values for a rectangle covering the edge (with a width equal to sigma) at an angle orthogonal to the edge for all four edges of the target (top, bottom, left and right). This allowed us to investigate whether the placement of patterning has an effect on fitness; for example, it has been suggested that stripes on the leading edge of a target may redirect capture attempts posteriorly[13].

The remaining data analysis was run in R (version 3.5.0) [34] and linear mixed models were fitted using lme4 (version 1.1-21) [35]. We expected many of the measures of patterning to be autocorrelated and therefore we reduced the number of variables by determining which were the best predictors of capture time using linear mixed modelling. For each metric, we created a model with the log of fitness (the average capture time) as the dependent variable. Generation was included as a second order fixed effect to account for non-independence in capture time between generations, and population (fast, medium or slow) was also included as a fixed effect. Replicate ID was included as a random effect. Model AIC values were compared to determine which metrics best predicted capture times, within different categories: for luminance metrics, this was the standard deviation of the luminance, a sigma value of 4 for vertical stripes, a sigma value of 2 for horizontal stripes, a sigma value of 2 for diagonal stripes (with both diagonal directions pooled together) and for edge metrics, a sigma value of 8 for the right hand edge. In all of these cases, the measure was a highly significant predictor of average fitness (p < 0.001 for all metrics). An example of the model structure used is as follows:

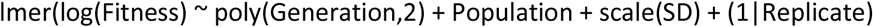

First, we modelled whether there was a change in fitness across generations and populations in our experimental data. We fit a model with the log of fitness as the dependent variable, and the second order effect of generation and the first order effect of population as fixed factors. Replicate number was included as a random slope. We then compared the change in fitness of our targets across generations for both the Eden project data and the null data, allowing us to test whether fitness improved in our experimental population compared to a null baseline. To do this, we fit a similar model as previously, but also included a variable indicating whether the data belonged to a null or an experimental population (’control’). The interaction between generation number and the ‘control’ variable was also included as the key interaction determining whether the increase in fitness was significantly different in the experimental population. Replicate ID was included as a random effect. This model also included only the first 40 generations.

We next tested whether there were differences in how our five patterning metrics had changed in the experimental and the null populations within the first 40 generations. To do this, we fit cumulative link models using the ordinal package (version 2019.4-25) [36], with generation as an ordinal dependent variable and the interaction between the metric and the ‘control’ variable as independent variables. We did not use the patterning metrics as dependent variables as these were highly skewed, making it difficult to fit an appropriate model, and we also did not use replicate ID as a random effect as this led to overfitting. The model included the first 40 generations. An example of the model structure used is as follows:

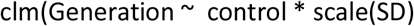

Finally, we wanted to analyse whether there were any differences in selection rates for the different speed populations in the experimental population over the first 40 generations. To do this, we used the Lande, Arnold and Wade framework [27–29] to calculate linear selection rates (β) for each of the five camouflage metrics within each population. For each combination of population, generation and replicate, we fitted a multiple linear regression between the dependent variable of logged fitness and the five normalised camouflage metrics as independent variables. Normalising the camouflage metrics ensured that the selection rates for each could be directly compared. We then took the linear regression coefficients for each metric as the linear selection rates. We used these to test for differences in linear selection rates between different speed populations and over evolutionary time (generations). We fitted linear mixed effect models using the linear regression coefficients for each metric as the dependent variable, testing against the second order fixed effect of generation and the fixed effect of population. Replicate ID was included as a random effect. An example of the model structure used was as follows:

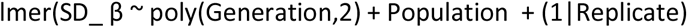

Significance tests for all models were carried out using the ‘Anova’ function from package ‘car’ (version 3.0-2) [37] which was used to calculate Type II ANOVAs. Where relevant, post-hoc comparisons were carried out with the ‘emmeans’ (version 1.3.4) package [38].

### Modelling methods

Motion modelling was carried out using a MATLAB implementation of a motion model using a two-dimensional array of correlation-type elementary motion detectors (as described in [39]) [40,41]. For each “fast” bug in generation 0 (512 bugs in total) we generated a short movie where the bug initially moved on an upwards trajectory and then rotated to move on a trajectory 15 degrees to the right (see supplementary material for an example). We used the generation 0 bugs as these should display a wide range of randomly selected pattern types, and the “fast” population as selection seemed to be strongest on these targets, suggesting that we should see the largest differences in fitness for this population. The time constant (tau) used was 3, the size of spacing between receptors was 50, the size of the filter was 30 and the standard deviations of the Gaussians (used for Difference of Gaussians spatial filtering) were 3 and 5.

For each bug, several metrics were calculated from the output of the motion model (after removing zeros, corresponding to places in the image where no motion signal was observed). Firstly, the mean resultant length of the circular direction data was calculated to give a measure of motion coherence. Secondly, the average vector length was calculated as a measure of motion energy. Finally, the bias was calculated by taking the difference between the circular mean and the “veridical” trajectory of the target (assumed to be the average of the two directions the target moved in during the trial). All circular statistics were calculated using CircStat [42].

Modelling was carried out using linear models, with the log of fitness being used as the dependent variable, and the coherence (mean resultant), bias (circular mean difference) and the motion energy (average vector length) were used as fixed factors in the model. The interaction between coherence and bias was also included, in line with predictions [19]. Finally, the data were filtered to include only the points with a circular mean difference of less than 60 degrees. The results were not qualitatively different if these data points were included. To test whether patterning metrics could predict the motion energy model output variables, we fit linear models with either the bias or the motion energy as independent variables, and either the standard deviation of the bug luminance or a metric of “stripy-ness”(the energy for vertical filtering angles with a sigma value of 4).

## Acknowledgements

AEH was supported by a PhD studentship from the BBSRC (BB/F016581/1) and is currently supported by a BBSRC grant (BB/P018319/1). LAK received funding from the People Programme (Marie Curie Actions) of the European Union’s Seventh Framework Programme (FP7/2007-2013) under REA grant agreement n° PIIF-GA-2012-327423 and is currently funded by a Royal Society Dorothy Hodgkin Fellowship. JT is funded by a NERC Independent Research Fellowship (NE/P018084/1). We would like to thank Amber Griffiths for providing valuable assistance and the Eden Project for hosting Dazzle Bug.

## Author contributions

AEH: conceptualisation, data curation, formal analysis, investigation, methodology, visualisation, writing (original draft preparation)

DG: data curation, formal analysis, investigation, methodology, software, writing (review and editing)

JT: data curation, formal analysis, software, writing (review and editing)

LAK: conceptualisation, funding acquisition, investigation, methodology, project administration, writing (review and editing)

